# High prevalence of *Klebsiella pneumoniae* in European food products: a multicentric study comparing culture and molecular detection methods

**DOI:** 10.1101/2021.11.24.469859

**Authors:** Carla Rodrigues, Kathrin Hauser, Niamh Cahill, Małgorzata Ligowska-Marzęta, Gabriella Centorotola, Alessandra Cornacchia, Raquel Garcia Fierro, Marisa Haenni, Eva Møller Nielsen, Pascal Piveteau, Elodie Barbier, Dearbháile Morris, Francesco Pomilio, Sylvain Brisse

**Affiliations:** Institut Pasteur, Université de Paris, Biodiversity and Epidemiology of Bacterial Pathogens, Paris, France; Institute for Medical Microbiology and Hygiene, Austrian Agency for Health and Food Safety, Vienna/Graz, Austria; Antimicrobial Resistance and Microbial Ecology Group, School of Medicine, National University of Ireland Galway, Ireland; Statens Serum Institut, Copenhagen, Denmark; Istituto Zooprofilattico Sperimentale dell’Abruzzo e del Molise “G. Caporale”, Teramo, Italy; Unité Antibiorésistance et Virulence Bactériennes, Université Claude Bernard Lyon 1 - ANSES, Lyon, France; INRAE, UR OPPALE, Rennes, France; Agroécologie, AgroSup Dijon, INRAE, Université Bourgogne Franche-Comté, Dijon, France

**Keywords:** *Klebsiella pneumoniae*, food sector, transmission, chicken meat, salads, antibiotic resistance, One Health, genomics, surveillance, culture methods

## Abstract

*Klebsiella pneumoniae* species complex (KpSC) is a leading cause of multidrug-resistant human infections. To better understand the potential contribution of food as a vehicle of KpSC, we conducted a multicentric study to define an optimal culture method for its recovery from food matrices, and to characterize food isolates phenotypically and genotypically. Chicken meat (n=160) and salad (n=145) samples were collected in five European countries and screened for KpSC presence using culture-based and ZKIR qPCR methods. Enrichment using buffered peptone water followed by streaking on Simmons citrate agar with inositol (44°C/48h) was defined as the most suitable selective culture method for KpSC recovery. High prevalence of KpSC was found in chicken meat (60% and 52% by ZKIR qPCR and culture approach, respectively) and salad (30% and 21%, respectively) samples. Genomic analyses revealed high genetic diversity with the dominance of phylogroups Kp1 (91%) and Kp3 (6%). 82% of isolates presented a natural antimicrobial susceptibility phenotype and genotype, with only four CTX-M-15-producing isolates detected. Notably, identical genotypes were found across samples: same food type and same country (15 cases); different food types and same country (1); same food type and two countries (1), suggesting high rates of transmission of KpSC within the food sector. Our study provides a novel isolation strategy for KpSC from food matrices and reinforces the view of food as a potential source of KpSC colonization in humans.

**Importance:** Bacteria of the *Klebsiella pneumoniae* species complex (KpSC) are ubiquitous and *K. pneumoniae* (Kp) is a leading cause of antibiotic-resistant infections in humans and animals. Despite the urgent public health threat represented by Kp, there is a lack of knowledge on the contribution of food sources to colonization and subsequent infection in humans. This is partly due to the absence of standardized methods for characterizing KpSC presence in food matrices. Our multicentric study provides and implements a novel isolation strategy for KpSC from food matrices and shows that KpSC members are highly prevalent in salads and chicken meat, reinforcing the view of food as a potential source of KpSC colonization in humans. Despite the large genetic diversity and the low-levels of resistance detected, the occurrence of identical genotypes across samples suggests high rates of transmission of KpSC within the food sector, which need to be further explored to define possible control strategies.

## Introduction

*Klebsiella pneumoniae* (Kp), a common gut bacteria, is regarded as a critical priority pathogen (1, 2), given the depletion of therapeutic options to treat multidrug-resistant (MDR) Kp infections. Furthermore, hypervirulent community-acquired invasive infections represent a growing problem, especially in Asian countries (3). Concerning reports of the convergence of MDR and hypervirulent phenotypes are increasing (4). The clinical epidemiology of Kp is dominated by the clonal spread of important MDR and hypervirulent sublineages, such as clonal group (CG) 258, CG15, CG147 or CG23 (3).

The ecology and transmission of Kp outside the clinical setting is still underexplored; in particular, the reservoir and possible contribution of non-clinical sources in the transmission of prominent clonal groups to humans remain largely undefined. Given the broad ecological distribution of Kp in animals and the environment (5, 6), and its capacity to contaminate food (7–9), the contribution of food sources to colonization of humans, and to subsequent infection, is an important question to address. Currently, detection, isolation and identification of Kp is not well integrated in food microbiological surveillance programs. There is also a lack of standardized tools and procedures for the detection of Kp in food. To define the natural ecology and transmission of Kp, studies of the presence of Kp in food should be designed irrespective of antimicrobial resistance phenotypes.

In order to achieve precise detection and identification of Kp, it is important to define the target species. Recent taxonomic updates pointed out the existence of seven phylogroups (phylogroup 1 [Kp1] to Kp7), now defined as distinct taxa, within *Klebsiella pneumoniae sensu lato*. The resulting *K. pneumoniae* species complex (KpSC) consists of five different species: *K. pneumoniae sensu stricto* (Kp1), *K. quasipneumoniae* subsp. *quasipneumoniae* (Kp2) and *K. quasipneumoniae* subsp. *similipneumoniae* (Kp4), *K. variicola* subsp. *variicola* (Kp3) and *K. variicola* subsp. *tropica* (Kp5), ‘*K. quasivariicola*’ (Kp6, which remains to be formally defined), and *K. africana* (Kp7) (10–12). Recently, Barbier *et. al* developed the ZKIR qPCR (13) for the detection of KpSC in soil and environmental samples, but its application to food matrices was not evaluated. Regarding the isolation of *Klebsiella* spp., different selective culture methods (7, 14, 15) have been described but comparisons of their efficiency for *Klebsiella* isolation from food is lacking.

The aims of this study were (i) To compare the selectivity, productivity and specificity of three agar media for the detection and isolation of *Klebsiella* spp., leading to propose a standardized culture protocol for the recovery of *Klebsiella* spp. in food matrices and compare its performance with the ZKIR qPCR; and (ii) To evaluate the presence and phenotypic and genomic features of KpSC in two common globally consumed foodstuffs: chicken meat and ready-to-eat salads, through a multicentric study in five European countries.

## Results

### Comparison of culture media

Productivity (P_R_), selectivity (S_F_) and specificity were calculated separately for each of the media considered. The three media were compliant with ISO11133:2014 requirements for productivity, as P_R_ was above 0.5 for almost all target strains **(Figure S3**). However, two strains had P_R_ < 0.50: *K. pneumoniae* SB132 in Chromatic Detection Agar and *K. quasipneumoniae* subsp. *quasipneumoniae* SB1124, for which colony growth was only observed on SCAI medium.

We observed that the media did not comply with the selectivity criteria (S_F_ > 2) for most of the non-target strains considered: the S_F_ values ranged from -0.3 to 0.5 for *Klebsiella* ChromoSelect Agar and from -0.5 to 0.9 for SCAI agar (Liofilchem was not tested as it is not selective). We noted that *Klebsiella* ChromoSelect Agar was selective for *Cronobacter* spp. and *C. freundii*, showing S_F_ values equal to 7.6 and 4, respectively. Hence, the media cannot be considered as selective for *Klebsiella* according to the ISO11133:2014 requirements.

Regarding specificity, the selective media tested showed variable results regarding the morphological features of observed colonies, based on the non-target strains considered. The selective media, indeed, allowed the growth of some non-*Klebsiella* spp. strains, and in few cases, colonies were morphologically similar to the target strains. Overall, because Kp colonies were easily distinguishable on SCAI medium and because it was as performant as the others regarding the other criteria, the SCAI medium was selected and used hereafter.

### Definition of an optimized protocol for recovery of KpSC from food samples

Our initial comparisons of the four different culture protocols (**Figure S1**) showed that the most effective method for recovery of KpSC from chicken meat was protocol 1B, which consists of a one-step enrichment in BPW. KpSC was recovered from 7 out of 36 (19.4%) samples using this protocol, while only 3 (8.3%) samples were positive using protocol 2 (enrichment in LB broth supplemented with ampicillin) and only 2 (5.6%) using protocol 1C (double enrichment: enrichment in BPW followed by enrichment in LB broth supplemented with ampicillin). No KpSC was isolated using direct plating (protocol 1A).

Further, using BPW enrichment of chicken meat (protocol 1B), a higher recovery of KpSC was observed when streaking a 10 µl loop onto SCAI agar plates, *versus* spreading 10 µl or 100 µl of bacterial enrichment. Of the 28 chicken meat samples examined using both methods, KpSC was recovered from 24 (85.7%) samples using the 10 µl loop *versus* 19 (79.2%) samples after spreading 10 µl, and 7 (29.2%) samples after spreading 100 µl, onto SCAI agar. Thus, protocol 1B combined with streaking was used henceforth.

### Evaluation of the SCAI agar plate incubation temperature

We compared two temperatures (37°C and 44°C) for the incubation of SCAI agar plates (**Figure S2**). Collectively, 111 chicken samples were tested and 67.7% (64/111) of them were positive for KpSC. KpSC was recovered from 59/111 (53%) samples when SCAI agar was incubated at 37°C, as well as when incubated at 44°C, indicating no difference between incubation temperatures (**Table S3**). However, when data from each individual institution was analyzed, a slightly higher percentage of positive samples was recorded following incubation of SCAI plates at 44°C at two institutions (AGES and NUIG), whereas contrasting results were observed at the third institution (SSI) (**Table S3**). As a result, a decision was made to implement incubation of SCAI agar plates at 44°C ± 1°C, which was incorporated into our protocol (**Figure 2**) (dx.doi.org/10.17504/protocols.io.baxtifnn). All further food samples were tested according to this protocol. Of note, no statistical differences were found in the prevalence of KpSC in from free-range and not free-range chicken meat samples (**Table S3**).

**Figure 1.**
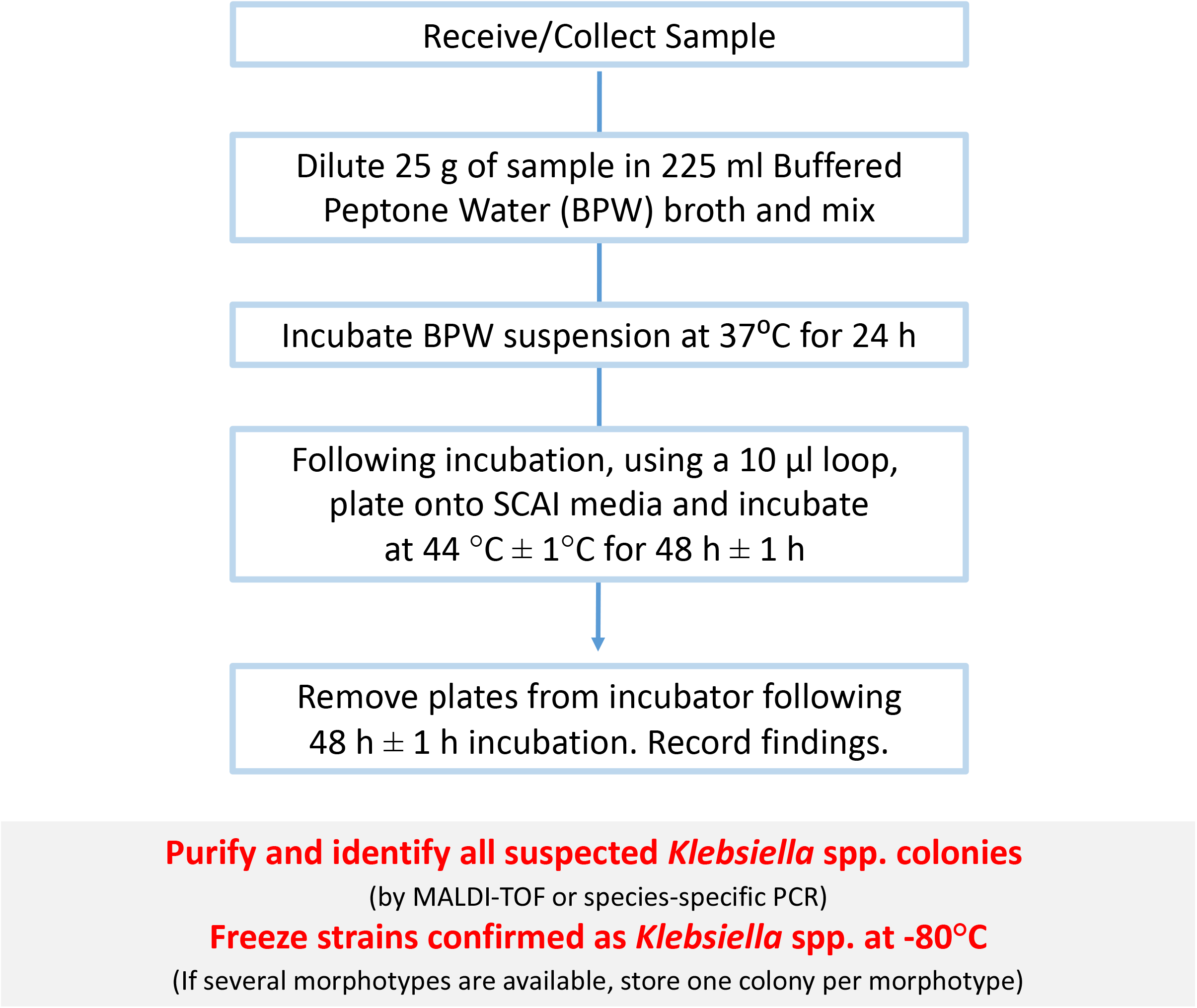
Optimized protocol for the recovery and isolation of *Klebsiella* spp. from food matrices.

**Figure 2.**
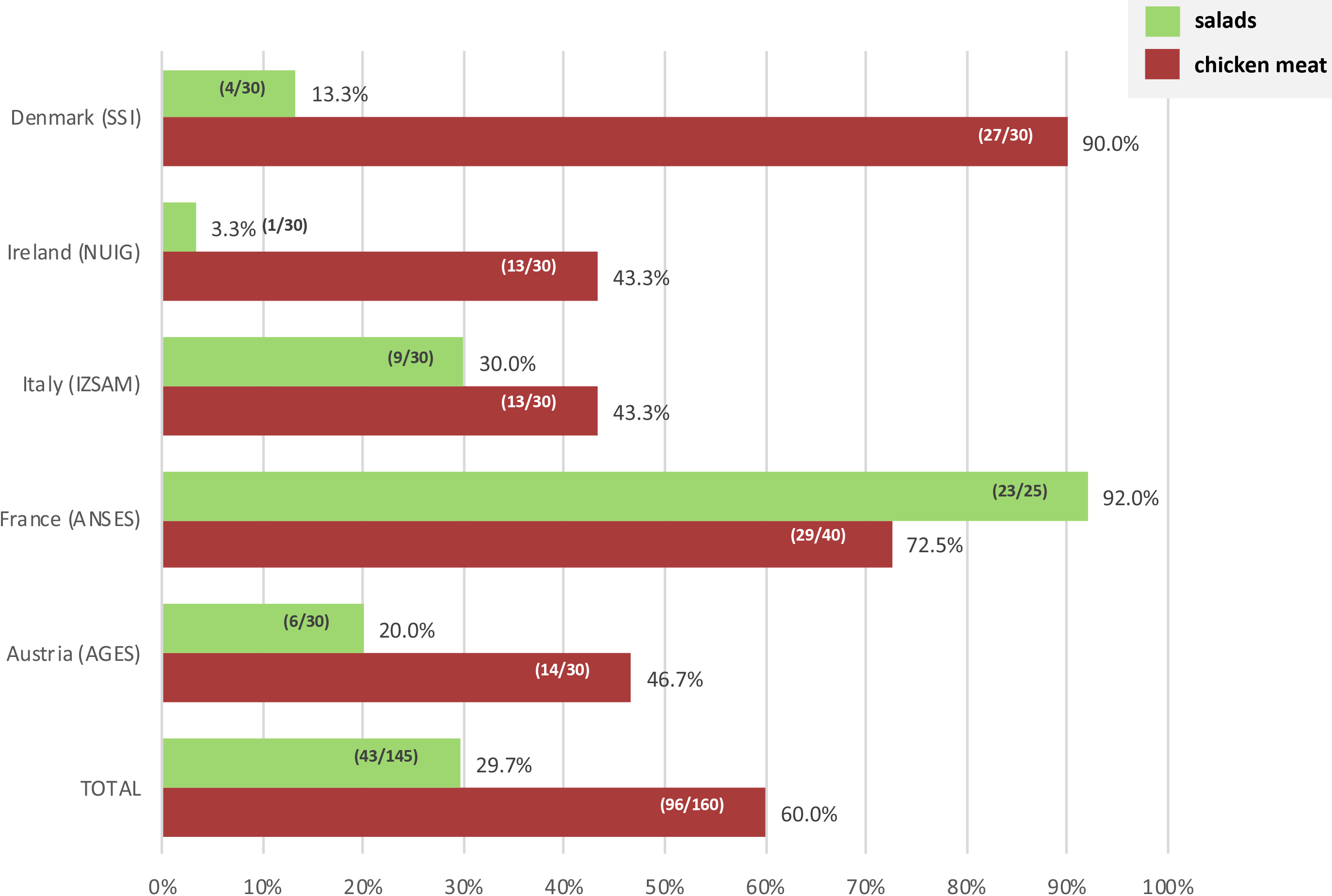
Prevalence of *K. pneumoniae* species complex by type of sample and by country.

### Comparison of culture and ZKIR qPCR methods for KpSC detection

Chicken meat and salad leaf samples were tested for the presence of KpSC using both the ZKIR qPCR assay (13) and the optimized culture protocol (**Figure 2**). Regarding the chicken meat samples (n=160), the ZKIR qPCR method detected KpSC in 96 (60.0%) of the samples (**Table 1**). In comparison, when the culture protocol was used, 83 (51.9%, *p*=0.177) of the samples were positive for KpSC (82 *K. pneumoniae* and 1 *K. variicola* according to MALDI-TOF MS results). Regarding salad leaf samples (n=145), 29.7% (43/145) of the samples tested positive for KpSC using the ZKIR qPCR assay, versus 20.7% (30/145; *p*=0.104; 18 *K. pneumoniae* and 12 *K. variicola* according to MALDI-TOF MS results) using the culture method (**Table 1**). Considering the ZKIR qPCR assay as a reference, the optimized culture protocol showed a sensitivity (true positive rate) of 84.2%, a specificity (false positive rate) of 100%, a positive predictive value of 100%, and a negative predictive value of 86.6%.

**Table 1.**
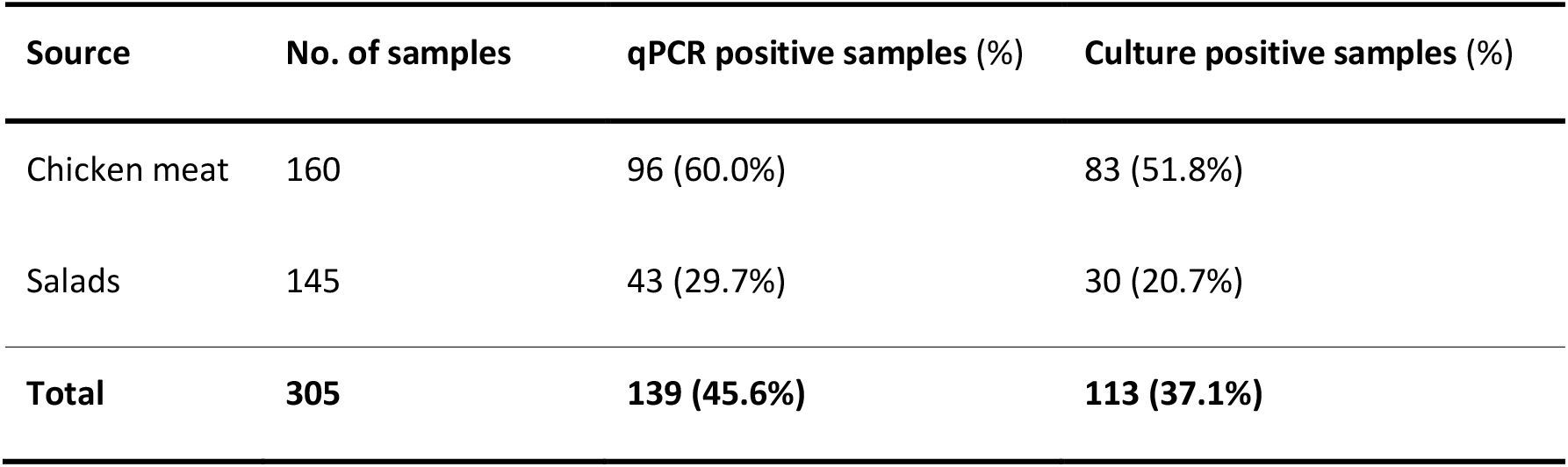
Comparison between the recovery rates of *K. pneumoniae* species complex using the ZKIR qPCR assay *versus* the optimized culture method.

| Source | No. of samples | qPCR positive samples (%) | Culture positive samples (%) |
| --- | --- | --- | --- |
| Chicken meat | 160 | 96 (60.0%) | 83 (51.8%) |
| Salads | 145 | 43 (29.7%) | 30 (20.7%) |
| **Total** | **305** | **139 (45.6%)** | **113 (37.1%)** |

### Comparison of KpSC prevalence in chicken meat and salad samples

The overall prevalence of KpSC in chicken meat was twice as high as in salad samples (60.0% versus 29.7%; *p* < 0.00001; **Table 1**). However, prevalence differed among countries (**Figure 2**). The prevalence of KpSC in chicken meat samples ranged between 43.3% and 46.6% for three of the countries (Austria, Italy and Ireland), whereas in France and Denmark the prevalence was above 70% (72.5% and 90.0%, respectively). Regarding salad samples, the prevalence of KpSC ranged from 3.3% in Ireland to 92% in France, with Austria, Italy and France being the countries where the highest numbers of samples positive for KpSC were detected.

In total, 131 KpSC isolates (92 from chicken meat and 39 from salads) were collected from 113 positive samples using the optimized culture method; in some positive samples, up to 5 different morphotypes were collected (**Figure 2, Table S2**).

### Antimicrobial susceptibility

Antimicrobial susceptibility phenotypes were defined for the 131 KpSC isolates using 17 antimicrobial agents representing 11 antimicrobial classes (**Table S4**). 82% of the isolates presented a wild-type phenotype, being susceptible to all the antibiotics tested except ampicillin, for which constitutive resistance is a characteristic of all KpSC isolates. The highest rates of resistance (detected in > 5% of the isolates) were observed for trimethoprim (12% in chicken meat and 13% in salads), trimethoprim-sulfamethoxazole (8% for both), tetracycline (9% in chicken meat and 5% in salads) and piperacillin-tazobactam (7% in chicken meat; **Figure S4**; **Table S4**). Four isolates (2 from each source) were found to be resistant to extended-spectrum cephalosporins, and ESBL gene presence in the genomic sequence was confirmed (**Table S4**). No isolate was found to be resistant to cefoxitin, ertapenem or netilmicin. Similar levels of resistance were detected among isolates from salads and from chicken meat.

Differences in antimicrobial resistance occurrence rates were observed among countries (**Table S4**) but the low number of isolates implies precaution in the analysis. For chicken meat, higher resistance rates were detected in Italy (8-50%), Austria (7-23%) and Ireland (7-14%), whereas for salads it was in Italy (33-100%), Austria (20%) and France (4-12%) (**Figure S4, Table S4**).

Nine (6.9%) multidrug resistant isolates were identified (**Table S5**), including the four ESBL producers. MDR isolates were recovered in 7 chicken meat and 2 salad samples in Italy (n=3), Austria (n=2), France (n=2) and Ireland (n=2) (**Table S5**).

### Phylogenetic and genomic diversity

Based on genomic data we observed among the KpSC isolates the predominance of Kp1 (90.8%, 119/131) followed by Kp3 (6.1%, 8/131), Kp2 (2.3%, 3/131) and Kp4 (0.8%, 1/131). The predominance of Kp1 was found in both sample types (98% in chicken meat; 74% in salads); however, there was an association of Kp3 with salad samples (18% in salads versus 1% in chicken meat; *p*=0.001). Kp2 was only found in salads, whereas Kp4 was only detected in chicken meat (**Table S6**).

Population structure analysis based on MLST and cgMLST genotyping unveiled a high genetic diversity among the KpSC isolates, with 86 STs [19 of them defined in this study; Simpson’s index of diversity (SID) 0.989] and 107 cgSTs (SID 0.995; **Figure 3**). High-risk clonal groups (CGs) common in the clinical setting represented 19.8% (26/131) of the isolates (**Table S6**). Among them, CG45 (n=9), CG37 (n=5), CG17 (n=4), CG661 (n=4), CG147 (n=2), CG14 (n=1) and CG15 (n=1) were found in chicken meat (n=18; 11.3%) and/or salad (n=6, 3.8%) samples. A high diversity of capsular types was also found, with 71 *wzi*-alleles and 58 KL-types. In contrast, O-type diversity was low, with O1 (25.1%; 33/131) and O2 (20.6%; 27/131) representing almost 50% of the recovered KpSC, followed by O3 (28.2%; 37/131) (**Figure 3**).

**Figure 3.**
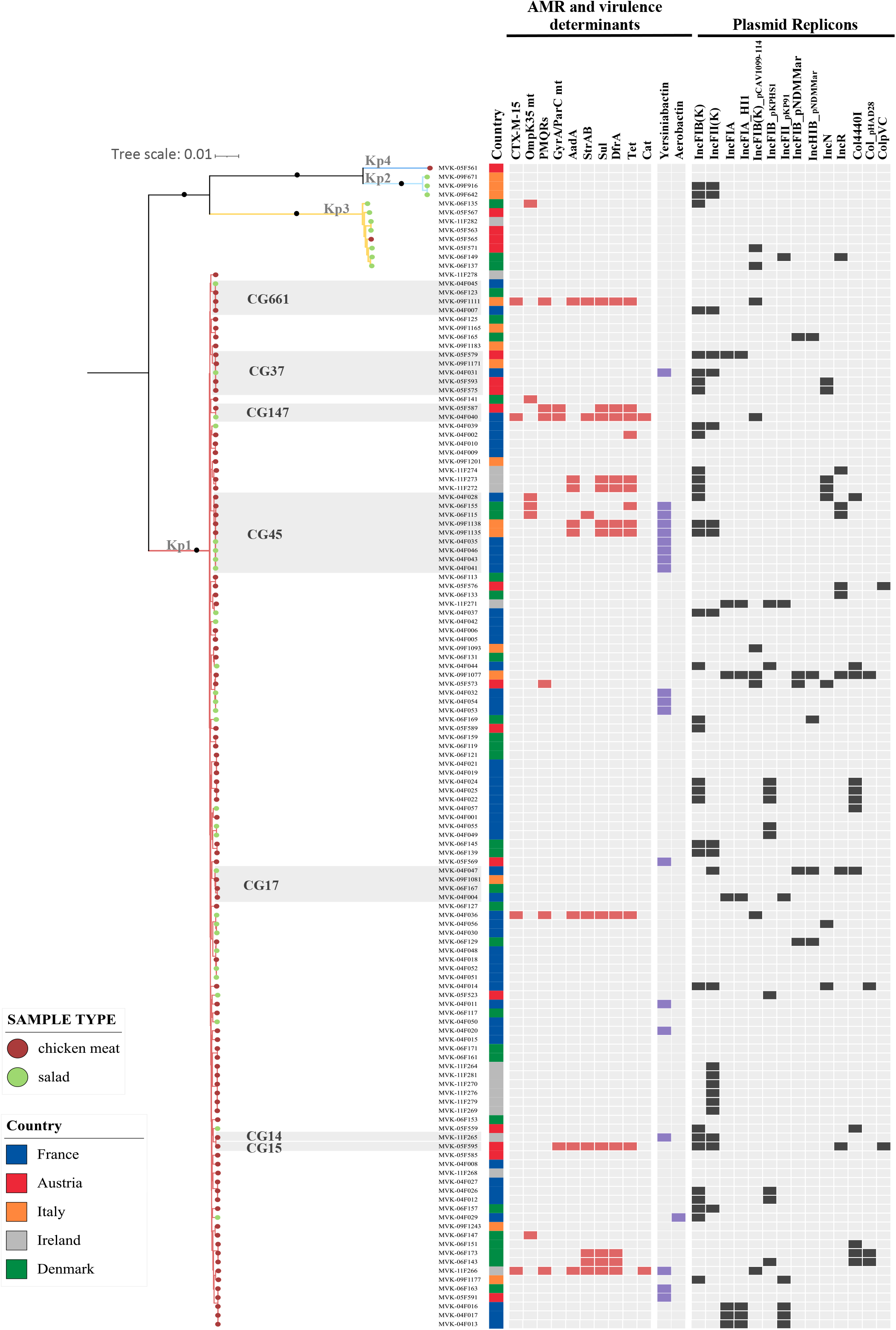
Maximum likelihood phylogeny based on 4,125 concatenated core genes generated using IQ-Tree with the GTR+F+ASC+R5 model. The scale bar indicates the number of nucleotide substitutions per site. Black dots on main nodes indicate bootstrap values ≥95%. Tree tips are colored by sample type (see key). The grey boxes delineate high-risk clonal groups (CG), as specified. European countries where the samples were collected are colored as indicated in the legend. The presence of antimicrobial resistance and virulence genes, and plasmid replicons is indicated.

Defining single strains (designed as genotypes) as isolates differing by no more than 5 cgMLST alleles (out of 629 loci) (16), we observed 15 strains that were isolated at least twice across a total of 35 different samples. These were collected from the same food type (chicken meat or salads) at different times in the same country (**Table S6, Figure S5**). To improve the confidence inferring the close relationship between these cgMLST-based overlapped genotypes, the SNP matrix based on 4,125 core genes obtained from Roary (representing ∼75% of a typical *K. pneumoniae* genome length against 10% represented by the 629 cgMLST genes) (16) was used, and confirmed the close genetic relatedness. For example, in France, the same genotype (ST45/genotype 7 – 1 to 4 SNPs) was detected in 3 salad samples from different brands (separated in time by 9 days), whereas in Ireland the same genotype (ST4384/genotype 13 – 1 to 8 SNPs) was detected in six different chicken meat samples (separated in time by 2 months) from different suppliers (**Table S6, Figure S5**). There was one case of the same strain being detected in different countries, as two Kp1 ST290 isolates (genotype 10; 15 SNPs) were detected in chicken meat samples from Denmark (April 2019) and Austria (September 2019) (**Figure S5**). We also detected a single strain between chicken meat and salads (ST1537/genotype 9 – 23 SNPs) between May and September 2019, respectively, in France (**Table S6**).

In most isolates (82%; 108/131) only intrinsic *bla*_SHV/OKP/LEN_ genes was detected, consistent with a natural susceptibility genotype (**Table S6**). Few antibiotic resistance genes were observed across the 131 isolates, with genes of non-prominent clinical significance *sul* and *dfr*, (9.2%; 12/131 both), *tet* (8.4%; 11/131), *aadA* and *strAB* (6.1%; 8/131 both) being the most frequent ones. However, four CTX-M-15 producing isolates were detected in two salad samples from France and two chicken meat samples from Italy and Ireland. The genotype of MDR isolates was compatible with their phenotype (**Table S5**). No carbapenemase or colistin resistance genes or mutations were detected. In 45.8% (60/131) of the isolates, neither plasmid replicons nor plasmid-encoded heavy metal tolerance genes were detected. In strains where plasmid replicons were detected, the plasmids known to occur naturally in Kp, such as IncFIB and IncFII_K_, were the most prevalent (25.2% and 16.8%, respectively). Regarding virulence genes, the yersiniabactin gene cluster was detected in 14.5% (19/131) of the isolates and no hypervirulent genotypes (as defined by the presence of *rmpA* or aerobactin or salmochelin) were detected.

## Discussion

Despite the renewed interest in *Klebsiella pneumoniae* complex (KpSC) epidemiology, boosted by the increasing involvement of *K. pneumoniae sensu stricto* (Kp1) in human infections associated with high-levels of antibiotic resistance and/or virulence (1, 3), the contribution of non-clinical sources, such as food products, to the current emergence of KpSC is poorly understood. This is partly due to the lack of standardized protocols for the detection and isolation of Kp strains in environmental, food or animal samples. Different selective culture media have been previously developed for *Klebsiella* spp.. Among the most recognized are MacConkey-Inositol-Carbenicillin agar (MCIC) (17) and its variation containing adonitol (MCICA,) (18), SCAI (14, 19) and Brilliant green containing Inositol-Nitrate-Deoxycholate agar (BIND) (20). All of them rely on the ability of *Klebsiella* spp. to use inositol as carbon source, combined with the use of secondary selective compounds (e.g., citrate as carbon source in the case of SCAI; nitrate as nitrogen source in BIND; carbenicillin and adonitol in MCIC/MCICA, although only Kp1 and Kp2/Kp4 are able to metabolize adonitol) (11). In our study, we decided to evaluate the performance of three different culture media for the detection and isolation of *Klebsiella* spp. in accordance with ISO 11133:2014 and latest amendments. They included the non-commercial SCAI medium and two commercial culture media: the *Klebsiella* ChromoSelect Selective Agar produced by Sigma-Aldrich, that similarly to MCIC uses carbenicillin as selective compound (https://www.sigmaaldrich.com/deepweb/assets/sigmaaldrich/product/documents/240/580/90925dat.pdf); and Chromatic Detection agar developed by Liofilchem, which is a chromogenic non-selective medium allowing the identification of *Klebsiella* spp., *Enterobacter* spp. and *Serratia* spp. based on the green-blue color of colonies (http://www.liofilchem.net/login/pd/ifu/11611_IFU.pdf). The productivity tests of the three media did not show any critical issues, whereas none of the media tested fulfilled the selectivity and specificity criteria defined by ISO 11133:2014. Nevertheless, the distinction of *Klebsiella* spp. colonies from other species was achieved in general. This is a critical point, as the routinely used media (e.g., MacConkey or CLED agar) do not provide easy differentiation of *Klebsiella* spp. colonies from other species, such as *Citrobacter* spp., *Pantoea* spp. or *Serratia* spp. Here, the performance of the tested culture media showed similar results and SCAI was selected, due to its low cost and easy in-house preparation. The use of this media in recent *Klebsiella* spp. epidemiological studies may also represent an advantage for global and cross-sector comparisons (13, 21–23).

Once SCAI agar was selected, different protocols to isolate *Klebsiella spp*. from food matrices were evaluated. Multiple conditions, involving pre-enrichment, enrichment, incubation temperature and plating were assessed from chicken meat samples. The optimized culture protocol was then compared with the reference ZKIR qPCR assay in a multicentric design study involving chicken meat and salad samples (13). Overall, KpSC was detected at a higher rate in both chicken meat and salad samples using the ZKIR qPCR method, but even so the optimized culture protocol demonstrated good sensitivity (84%) and specificity (100%) rates.

Approximately 50% of the food samples tested were KpSC positive, with significantly higher occurrence in chicken meat compared to salads, although disparities in prevalence were observed according to the country analyzed. It is difficult to contextualize our findings in a global scenario since few studies have addressed KpSC prevalence in these type of products without previous antibiotic selection: most of the studies addressing the prevalence and characterization of KpSC from foodstuffs have focused on colistin resistant or ESBL- or carbapenemase-producing *K. pneumoniae* (8, 24–26). As a consequence, knowledge on the ecology and natural diversity of Kp populations has been very limited. Furthermore, identification of the bacterial species was imprecise in most previous studies, in light of recent taxonomic changes. Our broader approach also captures susceptible strains and led to several important observations. First, the KpSC prevalence in chicken meat was higher than previously described: 47% in USA (Arizona) in 2012 (7); 30% in Turkey in 2007-2008 (27); 14% in China (Shijiazhuang) in 2013-2014 (28). Second, despite the high prevalence of KpSC in chicken meat samples, ESBL-producing *K. pneumoniae* were only detected in 1.3% (2 CTX-M-15) of the samples. These numbers are similar to those reported in other studies that targeted ESBL strains directly by selective approaches (0-5%) (8, 26, 28–30). Noteworthy, the presence of ESBL in chicken meat was scarce in *K. pneumoniae* when compared with *Escherichia coli* (55%-92%) in recent European reports (30–33).

Third, the highest resistance rates detected in natural food Kp populations were to tetracyclines, trimethoprim and trimethoprim-sulfamethoxazole, reflecting the recent reports of sales of these antimicrobial classes in food-producing animals in different European countries (34). Fourth, despite the predominance of Kp1 and the high genetic diversity found among chicken meat isolates, some genotypes overlapped across samples. The relation of these common genotypes (≤ 5 alleles mismatches) with the core-genome alignment obtained from Roary was also explored, and in most of the cases, less than 21 SNPs were identified, a threshold recently proposed for Kp ST258 to discriminate hospital clusters (35). Here, we found a maximum of 42 SNPs detected within the same genotype. Some of these multi-sample genotypes (ST290/genotype 10 recovered in Austria and Denmark and ST45/genotype 15 recovered in Denmark) were previously found circulating among healthy poultry in France in 2015 (Rodrigues C and Haenni M., pers.comm.). The repeated isolation of single strains in unrelated samples across time suggests transmission from the same source/reservoir or transmission within chicken flocks during production. Importantly, MDR ST45/genotype 11 found in Italy has also been found in urinary tract infections among Italian patients in 2018 (22), suggesting a contribution of food KpSC to human infections (7, 24).

Salads are typically eaten raw, with a high risk of ingestion of live Kp bacteria if present. The prevalence of KpSC among ready-to-eat salads was approximatively 30%, higher than what was reported in the few comparable studies (6-15%) (36–39). ESBL-producers were found in 1.4% of the samples (2 CTX-M-15), a lower prevalence than what is described in the literature (15-25%) targeting ESBL strains (39–42). As observed for chicken meat samples, the same genotype was observed within the same country, in salads from the same or different chain supplier/supermarket, suggesting a common source of the vegetables (e.g., different brands supplied by a common farm) or cross-contamination during their growth and processing (e.g., use of untreated irrigation water, wild-life animals, contamination of fields with manure, human contamination during packaging). Furthermore, one case of genotype overlap was found between salads and chicken meat isolates from samples from France, suggesting local transmission.

A previous study of risk factors of Kp carriage in community settings has linked the consumption of raw vegetables and contact with chicken to MDR KpSC carriage in humans (21). These findings together with our results, which are based on a small sampling survey, highlight the possible role of food as a source of Kp and call for much broader studies to address the ecology and transmission of this generalist pathogen in the food sector.

In our sampling, sublineages defined as high-risk clonal groups (3) represented 20% of the isolates recovered. One of these, ST45 (in both type of samples) was highly prevalent, as well as ST290 (in chicken meat), and the potential risk they pose should be addressed in future studies. Both STs have already been linked to human infections (22, 43, 44) and human carriage (21, 22) and also to broilers and pigs production (23, 45), raising the question of the zoonotic potential of these STs.

## Conclusions

In conclusion, our study addressed important knowledge gaps relating to Kp in food sources. First, we addressed the lack of harmonized culture protocols for the detection and isolation of *Klebsiella* spp. from food matrices, with the development of an optimized SCAI-based culture protocol, which in the future may be more widely adopted to enable global comparison regarding KpSC prevalence in these types of sources. Second, the high prevalence of KpSC in chicken meat and ready-to-eat salads highlights the possible role of food as a source of human colonization and infection by KpSC. Even though KpSC strains detected in food products remain largely susceptible to antimicrobials, understanding the degree to which food contamination by Kp contributes to human infections is an important topic for future research in the One Health perspective.

## Material and Methods

### Comparison of productivity, selectivity and specificity of agar media

Three different selective solid media for *Klebsiella* spp. growth were tested in accordance with ISO11133:2014 (Microbiology of food, animal feed and water — Preparation, production, storage and performance testing of culture media; https://www.iso.org/standard/53610.html). Nutrient agar (Microbiol & C., Cagliari, Italy) was used as the non-selective reference medium. For productivity, selectivity and specificity assays a set of 58 strains [50 from Institut Pasteur and 8 from Istituto Zooprofilattico Sperimentale dell’Abruzzo e del Molise Giuseppe Caporale (IZSAM)] were used: 51 reference strains of *Klebsiella* spp. and closely related species (*Raoultella* spp.), which included representatives of the six main KpSC phylogroups, and 7 non-target strains (**Table S1**). For productivity (P_R_), the three following media were tested: Simmons Citrate Agar with Inositol (SCAI) (14), *Klebsiella* ChromoSelect Selective agar produced by Sigma-Aldrich (Missouri, USA) and Chromatic Detection agar developed by Liofilchem (Roseto degli Abruzzi, Italy). The SCAI medium is not available commercially and was prepared in-house following previous protocols (14, 19) (https://www.protocols.io/view/isolation-of-klebsiella-strains-from-human-or-anim-662hhge/materials). P_R_ was calculated using 50 reference strains belonging to *Klebsiella* spp. and closely related species (*Raoultella* spp.) (**Table S1**). After incubation at 37°C for 24 h, broth cultures in Brain Heart Infusion (BHI) (Biolife, Milan, Italy) of each strain were diluted up to about 100 CFU/ml and spread on media. After incubation at 37°C for 48h, colonies compatible with the *Klebsiella* phenotype were enumerated. P_R_ is calculated as the ratio between the number of colonies grown on the selective agar medium and the number of colonies grown on the non-selective medium, and according to ISO11133:2014, P_R_ values should be ≥ 0.50 to consider the media productivity acceptable.

For selectivity (S_F_), 7 non-target strains (*Cronobacter* spp., *Citrobacter koseri, Citrobacter freundii*, *Serratia marcescens, Serratia liquefaciens, Serratia rubidaea* and *1 Pantoea agglomerans*) were tested on the SCAI and *Klebsiella* ChromoSelect Selective agar (**Table S1**). After incubation at 37°C for 24h in BHI, broth cultures were ten-fold diluted serially and each dilution was spread onto plates. After incubation at 37°C for 48 h, colony growth was observed. S_F_ is defined as the difference between the highest dilution showing growth on the non-selective reference medium (log_10_) and the highest dilution showing comparable growth on the selective test medium (log_10_). In accordance with ISO11133:2014, S_F_ values of non-target strains should be at least equal to 2 log_10_ growth differences, which reflects the ability of the medium to partially or totally inhibit their growth.

For specificity, the 7 non-target strains and two control strains (1 *K. pneumoniae* and 1 *R. ornithinolytica*) were tested on the three media (**Table S1**). Broth cultures were prepared in Buffered Peptone Water (BPW) (Biolife, Milan, Italy), incubated at 37°C for 24h, and spread onto each medium considered. After incubation at 37°C for 48h of each medium, plates were observed for the presence of typical and suspected colonies, recording if the characteristics of the non-target colonies could be similar to the target ones.

### Initial evaluation of protocols for recovery of *Klebsiella pneumoniae* from food

An initial assessment of four protocols for recovery of KpSC from chicken meat was carried out. In total, 36 chicken meat samples [12 free-range (7 skin-on and 5 skin-off) and 24 non-free range (12 skin-on and 12 skin-off) samples] were collected together with the following metadata: manufacturer, country of origin, and batch number. Each chicken meat sample was cut into small slivers to a final weight of 50 g. This 50 g sample was divided into two portions of 25 g and processed as follows and as illustrated in **Figure S1**:

**<u>Initial suspension 1</u>:** the first 25 g portion was diluted in 225 ml (1:10 dilution) of BPW and the sample was homogenized using a stomacher/blender. Prior to incubation of initial suspension 1, 100 µl of the suspension was cultured directly on SCAI medium using a sterile spreader or sterile loop and incubated at 37°C ± 1°C for 48h ± 1 h (**Figure S1 - 1A** - detection without enrichment). The remainder of initial suspension 1 was incubated at 37°C ± 1°C for 24h ± 1 h. Following 24h incubation of initial suspension 1, 100 µl of the suspension was streaked on SCAI medium and incubated at 37°C ± 1°C for 48 h ± 1 h (**Figure S1 - 1B** - detection with BPW enrichment only). Of the remaining incubated suspension 1, 1 ml was inoculated into 9 ml of Lysogeny broth (LB) with ampicillin (final concentration 0.01 mg/mL) and incubated at 37 °C ± 1 °C for 24h± 1 h. Following 24h incubation, 100 µl of the suspension was streaked on SCAI medium and incubated at 37°C ± 1°C for 48h ± 1 h (**Figure S1 - 1C** - detection with double enrichment).
**<u>Initial suspension 2</u>:** the second 25 g portion was diluted in 225 ml (1:10 dilution) of LB broth supplemented with ampicillin (final concentration 0.01 mg/mL) and the sample was homogenized using a stomacher/blender and incubated at 37 °C ± 1°C for 24 h ± 1 h. Following 24 h incubation, 100 µl of the suspension was cultured on SCAI medium and incubated at 37°C ± 1°C for 48 h ± 1 h (**Figure S1** - **2** detection with LB + Ampicillin enrichment only).

Typically, *Klebsiella* spp. appear yellow on SCAI medium (14). At least five suspected *Klebsiella* colonies were collected from each SCAI plate and sub-cultured onto SCAI media or a non-selective agar for identification using MALDI-TOF MS (Bruker Daltonics, Bremen, Germany). Currently, databases only allow for the identification of *K. pneumoniae* or *K. variicola*. Based on literature and in our test strain set, Kp1, Kp2 and Kp4 strains are identified as *K. pneumoniae*, whereas Kp3, Kp5 and Kp6 are currently identified as *K. variicola* (46, 47).

For 28/36 chicken meat samples, the impact on KpSC recovery of culturing using a 10 µl loop versus the spreading of 10 µl and 100 µl of enrichments onto SCAI media was also compared.

### Evaluation of the optimized protocol and definition of an optimal temperature of incubation

Following this initial assessment process, the optimized protocol as outlined below and as illustrated in **Figure S2** was evaluated for recovery of KpSC from chicken meat samples, using a total of 111 samples of chicken meat [51 free-range (25 skin-on and 26 skin-off) and 60 non-free range (30 skin-on and 30 skin-off)].

Each sample was cut into small slivers to a final weight of 25 g. This was added to 225 ml of BPW broth (1:10 dilution) and the sample was mixed using a stomacher or blender. The suspension was then incubated at 37°C ± 1°C for 24h ± 1 h. Following incubation, the suspension was cultured for single colonies on SCAI agar using a 10 µl loop and incubated at two different temperatures, 37°C ± 1°C and at 44°C ± 1°C for 48 h ± 1 h (**Figure S2**), in order to establish an optimal temperature of incubation of SCAI medium for recovery of Kp from food matrices.

### Comparison of the optimized SCAI culture protocol and ZKIR qPCR

A comparison of the optimized SCAI culture protocol and the ZKIR qPCR for the detection and isolation of KpSC in food matrices was carried out in 5 European institutions, including the Austrian Agency for Health and Food Safety (AGES), French Agency for Food, Environmental and Occupational Health & Safety (ANSES), National University of Ireland Galway (NUIG), Istituto Zooprofilattico Sperimentale dell’Abruzzo e del Molise Giuseppe Caporale (IZSAM) and the Statens Serum Institute (SSI, Denmark).

Each institution was asked to collect and test, where possible, an additional 30 samples of chicken meat, as well as 30 pre-packaged, pre-washed salad leaf samples (**Table S2**). In total, 160 chicken meat samples and 145 salad samples, collected between December 2018 and September 2019, were tested using the updated optimized protocol (dx.doi.org/10.17504/protocols.io.baxtifnn) for recovery of KpSC from food matrices (**Figure 1**).

The ZKIR qPCR was used to detect the presence of KpSC in food samples (13). After enrichment, DNA was prepared for ZKIR qPCR. 500 μl of the enrichment broth were centrifuged (5 min at 5800 × g), washed with sterile water, and resuspended in 500 μl sterile water before boiling for 10 min. qPCR conditions were as previously described (13) (dx.doi.org/10.17504/protocols.io.7n6hmhe).

### Antimicrobial susceptibility testing

Antimicrobial susceptibility testing was performed on 131 KpSC isolates using either disk diffusion or microbroth dilution [GN2F panels (Sensititre, ThermoFischer Scientific)] methods. Clinical breakpoints were used for interpretation according to EUCAST guidelines (https://www.eucast.org/fileadmin/src/media/PDFs/EUCAST_files/Breakpoint_tables/v_9.0_Breakpoint_Tables.pdf), except for kanamycin and streptomycin (48). *Escherichia coli* ATCC25922 was used as control strain. The panel of antimicrobials tested included: amoxicillin/clavulanic acid (20/10 µg), piperacillin-tazobactam (both using disks of 30/6 µg, or GN2F plates), cefpodoxime (10 µg, GN2F) or cefotaxime (5 µg), cefoxitin (30 µg, GN2F), aztreonam (30 µg), ertapenem (10 µg), amikacin (30 µg, GN2F), gentamicin (10 µg, GN2F), netilmicin (10 µg), tobramycin (10 µg, GN2F), kanamycin (10 µg), streptomycin (30 µg), ciprofloxacin (5 µg, GN2F) or ofloxacin (5 µg), nalidixic acid (30 µg), trimethoprim (5 µg), trimethoprim-sulfamethoxazole (23.75 + 1.25 µg, GN2F), and tetracycline (30 µg). The presence of multidrug resistant isolates, which are defined as those resistant to at least one agent in 3 different classes of antimicrobials (49), was analyzed.

### Whole-genome sequencing and comparative genomic analyses

Whole-genome sequencing was performed on the 131 Kp isolates. Genomic DNA libraries were prepared using the Nextera XT Library Prep Kit (Illumina, San Diego, USA) following the manufacturer’s protocol. Illumina sequencing was performed at the five partners: AGES and SSI using MiSeq (2 x 250 bp paired-end sequencing; n=18 and n=31 isolates respectively); ANSES and NUIG using NovaSeq 6000 (2 x 150 bp paired-end sequencing; n=52 and n=15, respectively); and IZSAM using NextSeq 500/550 (2 x 150 bp paired-end sequencing; n=15). Genomic assemblies were obtained using SPAdes v3.9 (50) or Velvet v1.2.10 (51) and were annotated using Prokka v1.12 (52).

Multilocus sequence typing (MLST) (53), core-genome MLST (cgMLST) and cgLIN codes (16) were determined using the BIGSdb-Kp database (https://bigsdb.pasteur.fr/klebsiella/klebsiella.html). This web tool and Kleborate (54) (https://github.com/katholt/Kleborate) were used to look for antimicrobial resistance, virulence, and heavy metal tolerance genes and to characterize the capsular synthesis gene cluster. Plasmid replicons were searched using PlasmidFinder (55) (https://cge.cbs.dtu.dk/services/PlasmidFinder/).

For phylogenetic analyses, a core-genome alignment based on the concatenation of 4,125 core genes was obtained with Roary v3.12 (56) using a blastP identity cut-off of 80% and core genes defined as those being present in more than 90% of the isolates. Recombination events were removed using Gubbins v2.2.0 (57), generating a recombination-free alignment comprising 486,290 single-nucleotide variants (SNVs). This recombination-free alignment was used to construct a maximum likelihood phylogenetic tree using IQ-TREE v1.6.11 (model GTR+F+ASC+R5).

Chi square or Fisher exact test were used to compare the prevalence of KpSC in the different sources and to check the association of the different categorical variables (*P* values <0.05 were considered statistically significant).

### Data availability

Detailed optimized SCAI culture protocol for the detection and isolation of KpSC in food matrices was made publicly accessible to the scientific community in January 2020 through the protocols.io platform (dx.doi.org/10.17504/protocols.io.baxtifnn). Genomic sequences generated in this study were submitted to the European Nucleotide Archive and are accessible under the BioProject number PRJEB34643, and are also publicly available in BIGSdb through project ID 37 ‘MedVetKlebs_multicentric study’.

## Authors Contribution

SB, CR, DM, KH, EMN and FP conceptualized the study. GC, AC and FP from IZSAM performed the culture media evaluation. KH, MLM, EMN, NC and DM performed the initial assessments for the establishment of an optimized culture protocol for recovery of KpSC from food matrices. KH, NC, MLM, GB, AC and RGF performed the comparison of culture media and ZKIR qPCR assay. EB and PP provided laboratory support and validated the analysis of the ZKIR qPCR assay. CR ad SB analyzed all the genomic data, coordinated the study and wrote the initial draft manuscript. SB acquired funding. All authors contributed to writing and editing the manuscript and reviewed the final version.

## Acknowledgements

We thank to James Bray and Keith Jolley for technical assistance with genomic assemblies of KpSC isolates from Ireland. We thank the Institut Pasteur teams for the curation and maintenance of BIGSdb-Pasteur databases at http://bigsdb.pasteur.fr/.

## Authors license statement

This research was funded, in whole or in part, by Institut Pasteur and by European Union’s Horizon 2020 research and innovation programme. For the purpose of open access, the authors have applied a CC-BY public copyright license to any Author Manuscript version arising from this submission.

## Supplementary Material

**Table S1**. Bacterial reference strains used for agar media performance comparison.

Strains used for the different evaluation criteria are indicated. The correspondent strain code for productivity results in Figure S3 is indicated.

**Table S2**. Chicken meat and salad samples analyzed in the different European countries for the presence of *K. pneumoniae* species complex and its epidemiologic characteristics.

**Table S3**. Comparison between the recovery rates of *K. pneumoniae* species complex using two different incubation temperatures of SCAI medium.

**Table S4**. Number of *K. pneumoniae* species complex resistant isolates for each of the antimicrobials tested distributed by sampling source and by country.

**Table S5**. Multidrug resistant strains from chicken meat and salads, sequence-type (ST), phenotypic resistance and identified AMR genes based on WGS.

**Table S6.** Characteristics of the 131 *K. pneumoniae* species complex isolates recovered: population structure, surface polysaccharides, antimicrobial susceptibility, antimicrobial resistance genes, virulence genes, metal tolerance gene clusters and plasmid replicons.

**Figure S1.** Evaluation of different protocols for recovery of *Klebsiella* spp. from food matrices.

**Figure S2.** Protocol to define the optimal temperature of incubation of SCAI medium plates for the recovery of *Klebsiella* spp. from food matrices.

**Figure S3.** Productivity (P_R_) results for the three media considered using a reference panel of 50 *Klebsiella* spp. and closely related species.

**Figure S4.** Antibiotic resistance of the *K. pneumoniae* species complex isolates recovered by sample type and by country.

Upper panel: The x-axis represents the number of KpSC resistant isolates for each of the antimicrobials tested and percentages (%) displayed on the graph were calculated based on the total number of isolates.

Lower panel: The x-axis represents the number of KpSC resistant isolates for each of the antimicrobials tested and percentages (%) displayed on the graph were calculated based on the total number of isolates in each country.

**Figure S5.** Geographic distribution of the common genotypes found among *K. pneumoniae* species complex recovered from food products.

