## Supplementary material for "High prevalence of *Klebsiella pneumoniae* in European food products: a multicentric study comparing culture and molecular detection methods": Figure S1 to S5

**Figure S1** – Evaluation of different protocols for recovery of *Klebsiella* spp. from food matrices.

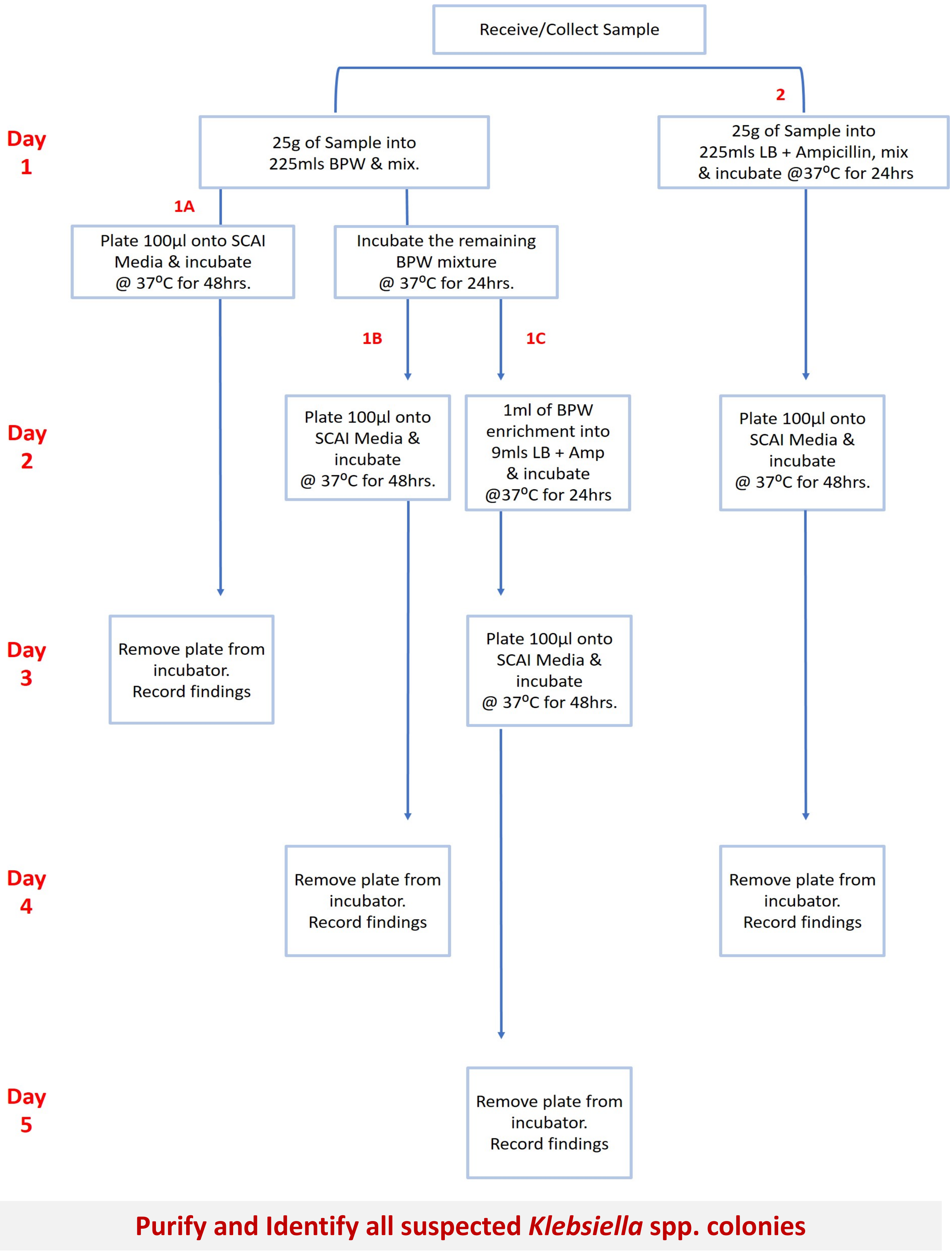


BPW, Buffered Peptone Water
SCAI, Simmons Citrate Agar with Inositol
LB, Lysogeny broth
Amp, Ampicillin
@, at

**Figure S2** - Protocol to define the optimal temperature of incubation of SCAI medium plates for the recovery of *Klebsiella* spp. from food matrices.

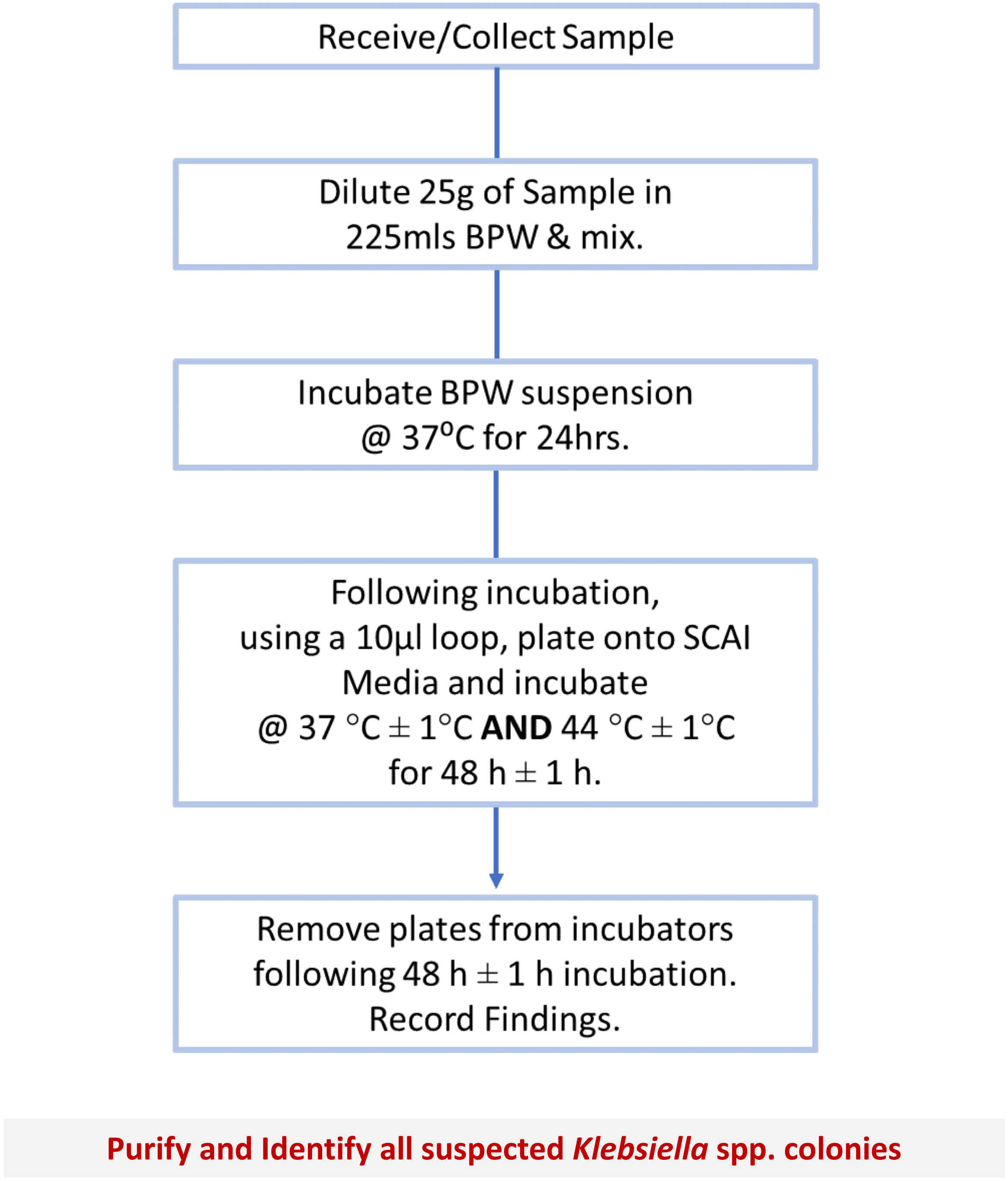


BPW, Buffered Peptone Water
SCAI, Simmons Citrate Agar with Inositol
@, at

**Figure S3**. Productivity ($P_R$) results for the three media considered using a reference panel of 50 *Klebsiella* spp. and closely related species.

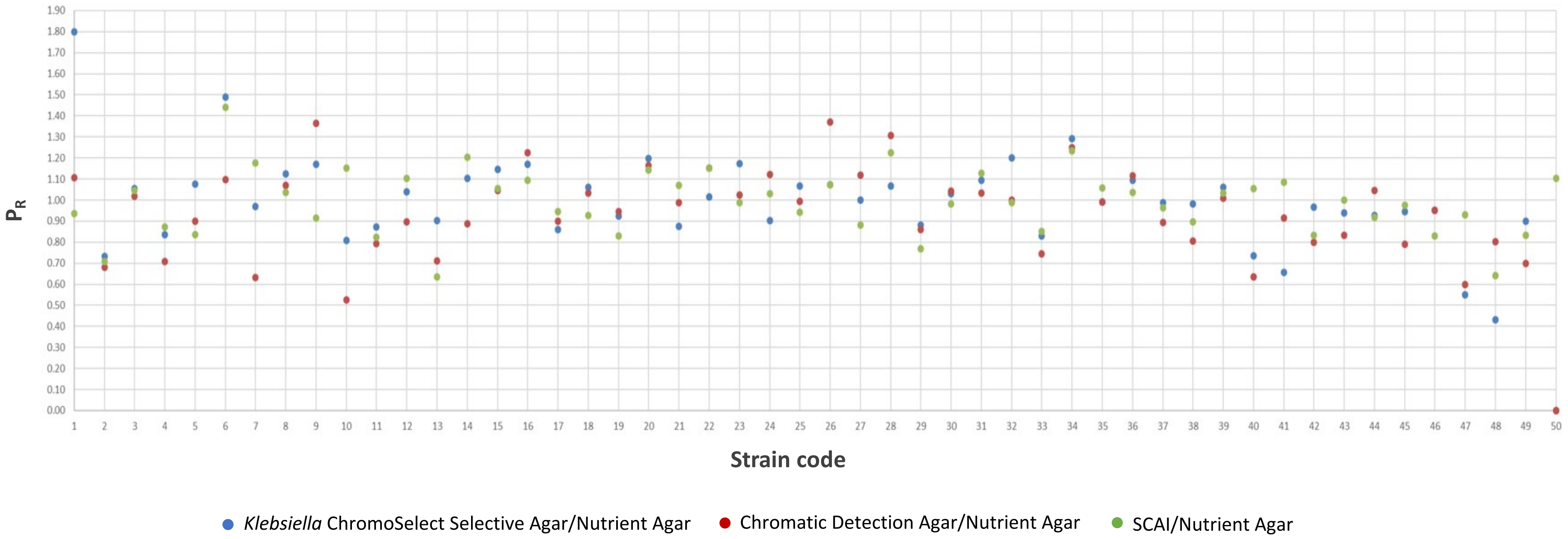

**Figure S4**. Antibiotic resistance of the *K. pneumoniae* species complex isolates recovered by sample type and by partner.

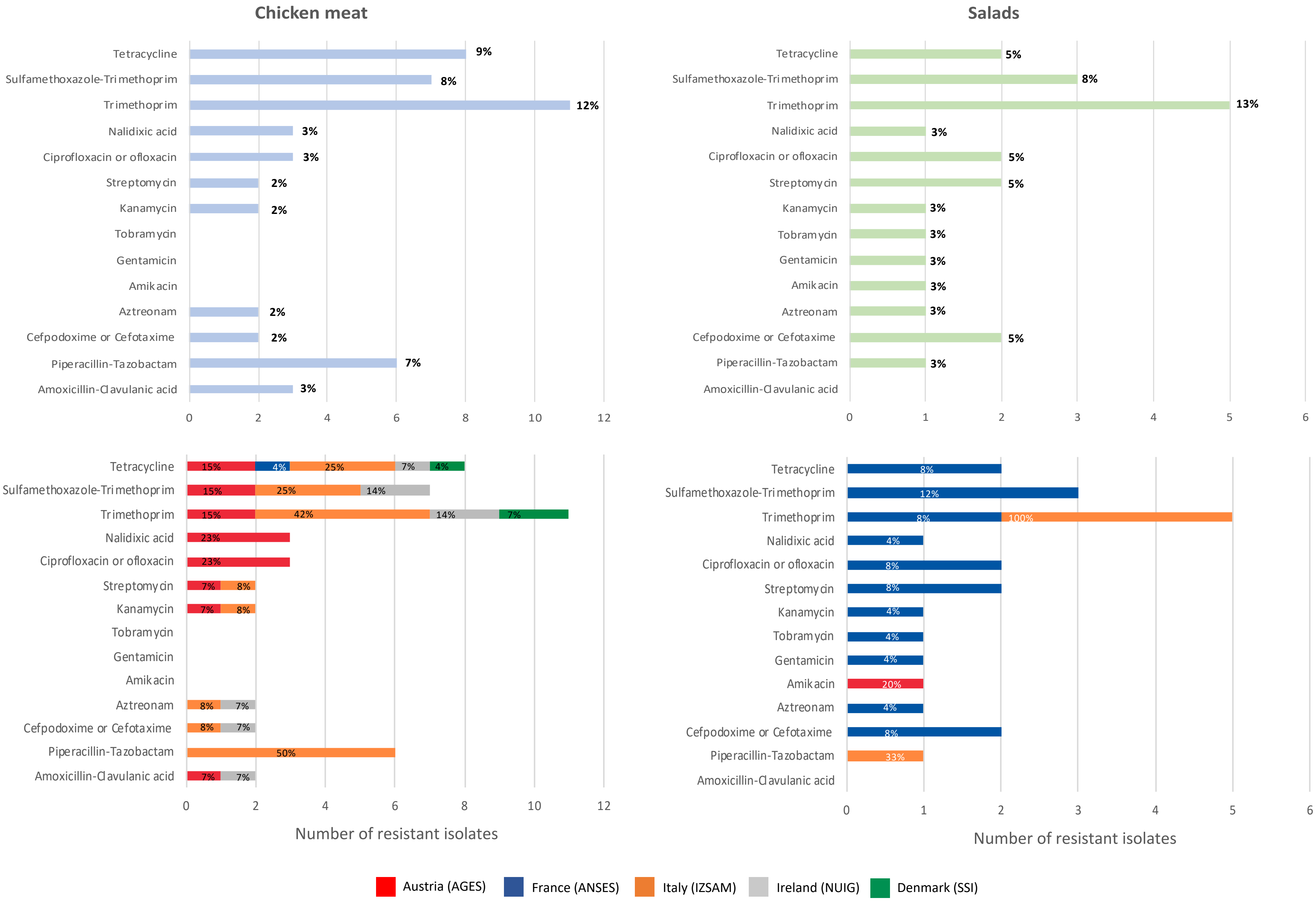

**Figure S5**. Geographic distribution of the common genotypes found among *K. pneumoniae* species complex recovered from food products.

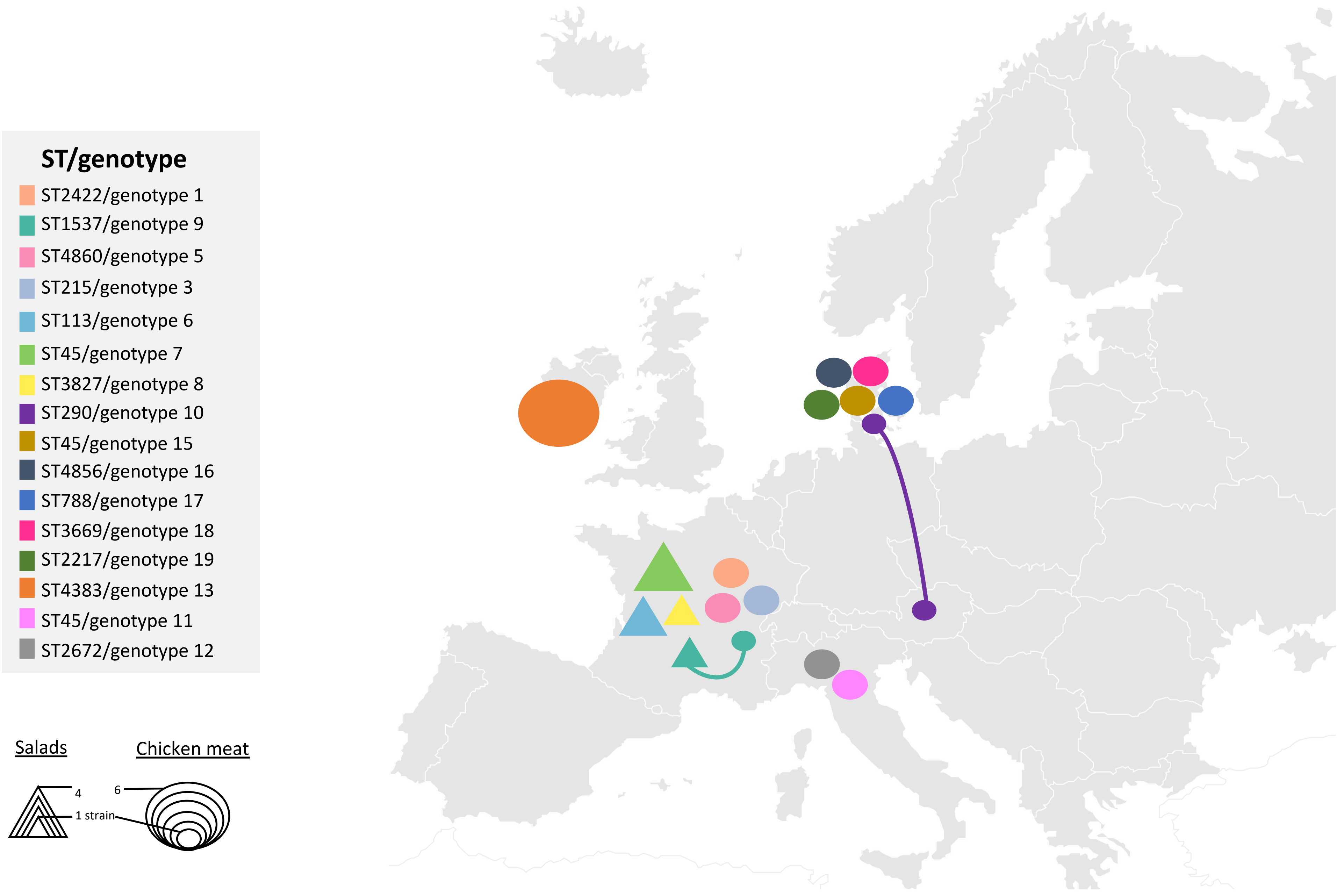
